# Mineralocorticoid receptor-independent activation of ENaC in bile duct ligated mice

**DOI:** 10.1101/2023.09.19.558474

**Authors:** Xue-Ping Wang, Stephanie M. Mutchler, Rolando Carrisoza-Gáytan, Mohammad Al-Bataineh, Catherine J. Baty, Amber Vandevender, Priyanka Srinivasan, Roderick J. Tan, Michael J. Jurczak, Lisa M. Satlin, Ossama B. Kashlan

**Author notes:** Correspondence: Ossama Kashlan, Department of Medicine, Renal-Electrolyte Division, University of Pittsburgh, S828B Scaife Hall, 3550 Terrace St., Pittsburgh, Pennsylvania 15261. These authors contributed equally (Xue-Ping Wang, Stephanie M. Mutchler).

## Abstract

Sodium and fluid retention in liver disease is classically thought to result from reduced effective circulating volume and stimulation of the renin-angiotensin-aldosterone system (RAAS). Aldosterone dives Na^+^ retention by activating the mineralocorticoid receptor and promoting the maturation and apical surface expression of the epithelial Na^+^ channel (ENaC), found in the aldosterone-sensitive distal nephron. However, evidence of fluid retention without RAAS activation suggests the involvement of additional mechanisms. Liver disease can greatly increase plasma and urinary bile acid concentrations and have been shown to activate ENaC *in vitro*. We hypothesize that elevated bile acids in liver disease activate ENaC and drive fluid retention independent of RAAS. We therefore increased circulating bile acids in mice through bile duct ligation (BDL) and measured effects on urine and body composition, while using spironolactone to antagonize the mineralocorticoid receptor. We found BDL lowered blood [K^+^] and hematocrit, and increased benzamil-sensitive natriuresis compared to sham, consistent with ENaC activation. BDL mice also gained significantly more body water. Blocking ENaC reversed fluid gains in BDL mice but had no effect in shams. In isolated collecting ducts from rabbits, taurocholic acid stimulated net Na^+^ absorption but had no effect on K^+^ secretion or flow-dependent ion fluxes. Our results provide experimental evidence for a novel aldosterone-independent mechanism for sodium and fluid retention in liver disease which may provide additional therapeutic options for liver disease patients.

**Significance:** Advanced liver disease is often complicated by renal sodium retention, leading to fluid retention, poor outcomes, and increased mortality. This is currently thought to be driven by increased levels of the hormone aldosterone, although numerous published reports demonstrate normal aldosterone levels in many liver patients with volume overload. Management of these patients relies on diuretics and Na^+^ restriction, or more invasive procedures with increased risks for diuretic-resistant patients. Here, we report a novel mechanism for fluid retention in liver disease, which provides a basis to develop new strategies to treat fluid retention in these patients.

## Introduction

Advanced liver disease often features Na^+^ retention, contributing to ascites, edema, and electrolyte imbalances. According to accepted paradigms, portal hypertension with peripheral vasodilation leads to a deficit in the effective circulating volume, which stimulates the renin-angiotensin-aldosterone system (RAAS) to promote renal Na^+^ and fluid retention and ultimately restore volume.^1^ However, numerous studies over the past 40 years document that roughly half of examined liver disease patients exhibiting fluid retention did not have RAAS activation.^2-11^ This suggests that a deficit in effective volume is insufficient to explain fluid retention in these patients.

Some liver conditions are also characterized by greatly elevated concentrations of plasma and urinary bile acids.^12-16^ Poor bile flow (i.e., cholestasis) reduces the conversion of primary bile acids to secondary bile acids by gut microbes. Ultimately, cholestasis induces spillover of chiefly conjugated primary bile acids to the systemic circulation.^16,17^ We and others reported that conjugated primary bile acids activate the epithelial Na^+^ channel (ENaC), a major target of aldosterone, *in vitro*.^18-21^ These compounds directly bind the channel and increase the likelihood of the channel being open.^19^ In principal cells of the connecting tubule and collecting duct, ENaC mediates the rate-limiting step for Na^+^ reabsorption.^22^ Electrogenic ENaC-mediated Na^+^ reabsorption provides the driving force for K^+^ secretion into urine through apical K^+^ channels, contributing to bodily K^+^ loss.^23^

In liver disease, activation of ENaC is evidenced by the important role of MR antagonists (e.g., spironolactone) in treating fluid retention.^1^ As urine becomes a major vehicle for bile acid excretion in advanced liver diseases,^13,24^ bile acids may activate ENaC independent of MR signaling. Here, we tested that hypothesis by increasing bile acids through bile duct ligation (BDL) and examining effects on renal function and body composition. With spironolactone present to inhibit MR, BDL increased ENaC-dependent Na^+^ retention and rendered total body water highly responsive to ENaC blockers. Changes in transporter expression or localization did not account for these effects on ENaC function. In isolated rabbit cortical collecting ducts, taurocholic acid (t-CA) increased net transepithelial Na^+^ absorption but had no effect on K^+^ secretion. Additionally, flow-dependent stimulation of Na^+^ absorption and K^+^ secretion was preserved in the presence of t-CA. Our data provide evidence for the activation of ENaC independent of MR signaling in a cholestatic liver disease model.

## Materials and methods

The materials and methods in this study are available in the supplementary materials.

## Results

### BDL elevates plasma bile acids and aldosterone

To examine the effect of elevated bile acids on renal function *in vivo*, we performed BDL and sham surgeries on 8-12 week old mice and sacrificed them 11 days later to collect blood and tissues for analysis. BDL resulted in jaundice and darkened urines, and increased plasma bile acids 50-fold versus sham operation to 1.6 ± 0.7 mM (Figure 1A). Plasma aldosterone levels were also higher after BDL (Figure 1B), as previously reported.^25^ Consistent with this, BDL mice had lower blood K^+^ and metabolic alkalosis, indicated by higher tCO_2_ and lower anion gap values (Table 1). BDL mice also had 10% lower hematocrit, which may reflect fluid retention and/or eryptosis.^26,27^ Kidney sections revealed tubular injury manifesting as dilated tubules and protein casts in BDL mice (Figure 2A, B). BDL also increased transcript levels for the kidney injury markers KIM-1 and NGAL by 3.5-fold and 43-fold, respectively, compared to sham (Figure 2C).^28,29^

**Table 1.**
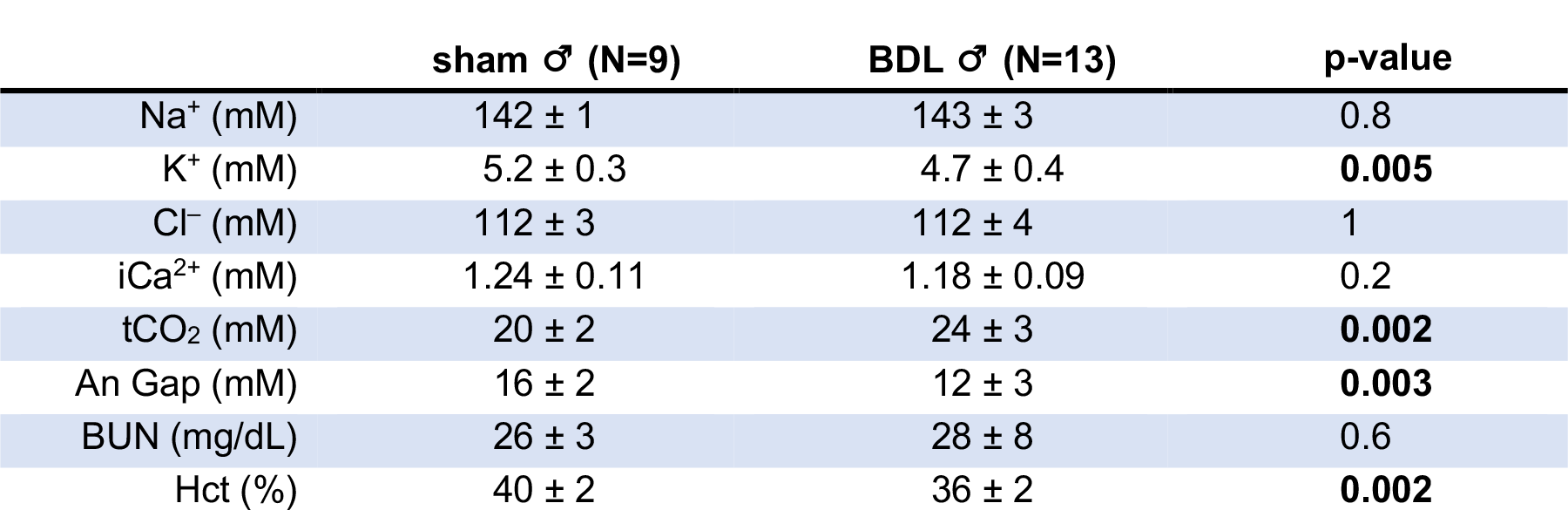
Blood parameters for sham and BDL mice. Blood was collected 11 days after surgery. Groups were compared by Student’s *t* test. The number of mice in each group and the mean ± SD for each parameter are shown. Blood creatinine was < 0.2 mg/dL for all mice, except for 1 BDL mouse (0.3 mg/dL).

**Figure 1.**
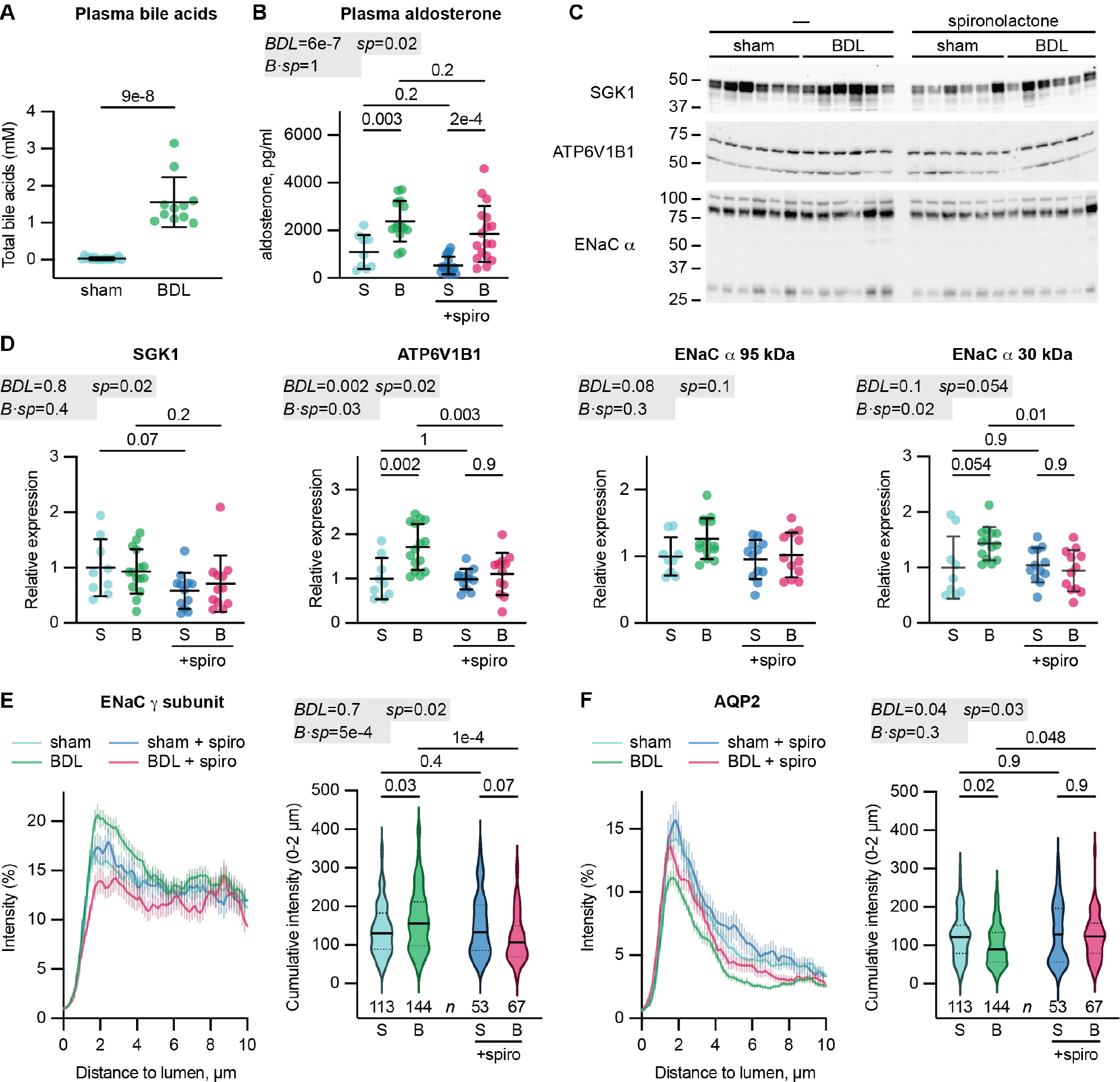
BDL increases bile acids and aldosterone, and spironolactone attenuates aldosteronism effects. The effect of BDL in the presence or absence of spironolactone (spiro) was determined. *A*, Total plasma bile acids after BDL and sham surgeries (Mann-Whitney U test). *B*, Plasma aldosterone after BDL (B) and sham (S) surgeries in the presence or absence of spironolactone (spiro) treatment. *C–D*, The effect of surgery and treatment on expression of indicated proteins was analyzed by immunoblot, and quantified and normalized against the stain-free gel. Values were normalized to the mean of untreated sham mice. Data for female mice treated with spironolactone are in Supplemental Figure 1. Individual values with mean and SD are shown in *A–D*. Data in *B–F* were compared by two-way ANOVA (shaded gray). *E–F*, Line scans on maximum projection images were drawn over ENaC and AQP2 positive cells and aligned to their apical edges. Twelve images from the outer cortex, middle cortex and juxtamedullary regions in each of 3 animals from each group were evaluated. The mean intensity profile ± SEM for each group is shown. The cumulative signal intensity within 2 μm of the apical edge was compared by two-way ANOVA (shaded gray). Data are shown as violin plots with median (solid line) and quartiles (dotted lines) indicated.

**Figure 2.**
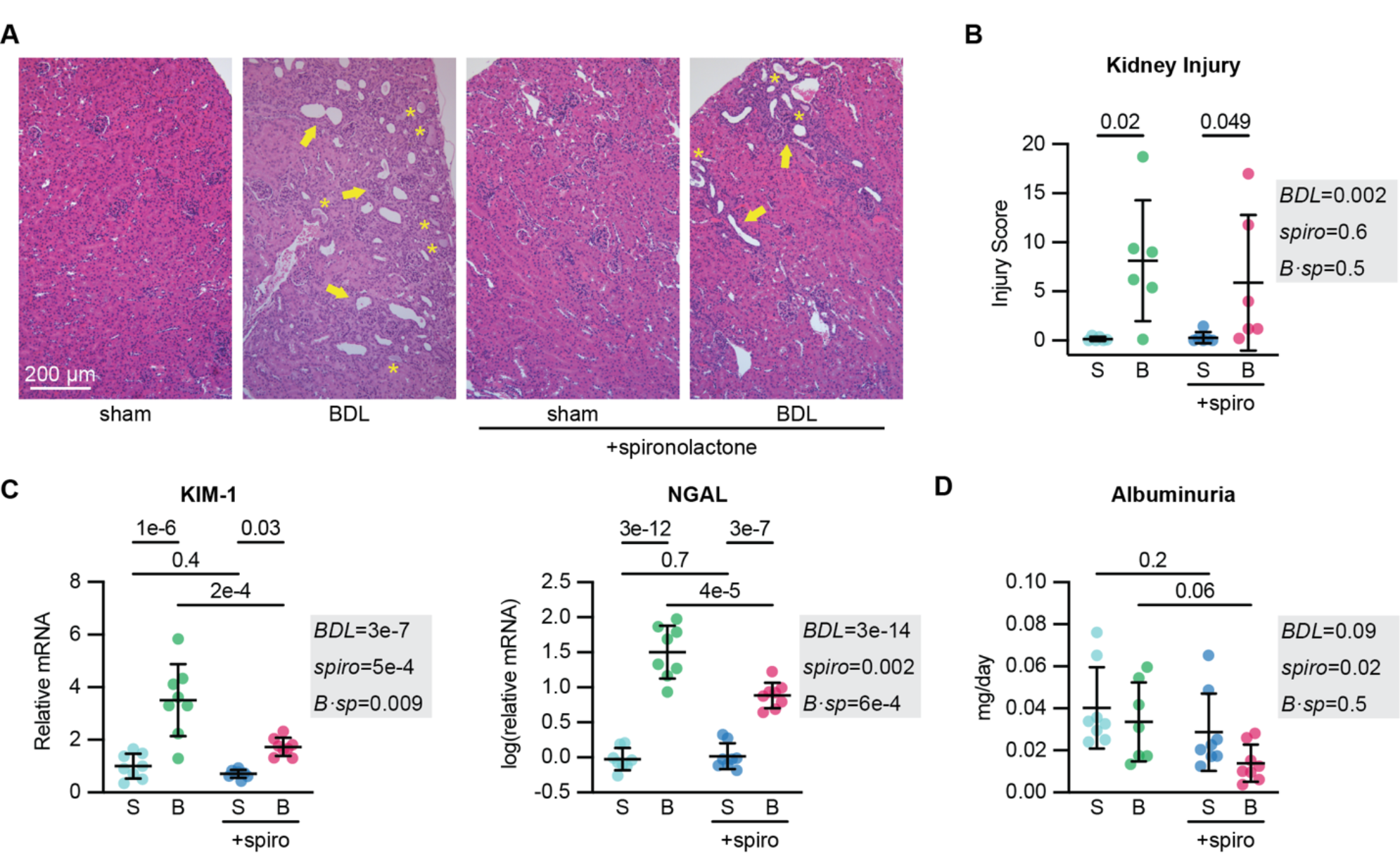
BDL causes kidney injury. *A*, Hematoxylin and eosin-stained kidney sections, with arrows indicating dilated tubules and regions of inflammation, and asterisks indicating casts. *B*, Blinded injury scores of kidney sections. *C*, Transcript levels of kidney injury markers NGAL and KIM-1 were measured by qPCR from kidney homogenates. Values were normalized to the mean of untreated sham mice, and NGAL values were log-transformed prior to comparison. *D*, Urinary albumin was measured from a 24-hour collection 7 days after surgery. Groups were compared by two-way ANOVA (shaded gray).

### Spironolactone normalizes secondary effects of aldosterone after BDL

To separate the effects of BDL on renal function from the secondary effects of MR activation by aldosterone, we began supplementing the drinking water on day 3 for both groups with spironolactone (20 mg·kg^-1^·day^-1^),^30^ an MR antagonist. BDL mice continued to have elevated aldosterone (Figure 1B, Supplemental Figure 1) and reduced hematocrit (Figure 3 and Supplemental Table 2), however, spironolactone normalized some of the effects of BDL that could be attributed to the secondary effects of aldosterone acting through MR. First, spironolactone rescued the metabolic alkalosis seen in BDL animals. Although BDL mildly increased overall tCO_2_ levels in spironolactone treated animals (Supplemental Table 2), we detected no differences between sham and BDL in males or females (Figure 3). Anion gap values were also similar between sham and BDL mice. Aldosterone increases the expression of several proteins, including the B1 subunit of the vacuolar ATPase proton pump (ATP6V1B1),^31,32^ the serum- and glucocorticoid-regulated kinase (SGK1),^33^ and the α subunit of ENaC.^34^Consistent with this, BDL increased expression of ATP6V1B1, while spironolactone treatment reduced expression to levels seen in shams (Figure 1C-D, Supplemental Figure 1). BDL did not affect SGK1 expression, although spironolactone had a mildly dampening effect (Figure 1C-D, Supplemental Figure 1). Any effect of BDL on ENaC α subunit expression was below detection; however, spironolactone did reduce expression of the cleaved 30 kDa form associated with activation. Furthermore, in immunofluorescence-stained kidney sections, spironolactone reversed BDL-induced changes in the localization of ENaC *γ* subunit and aquaporin-2 (AQP2). In the absence of spironolactone, BDL mice had greater ENaC *γ* subunit immunofluorescence intensity and lower AQP2 intensity near the apical surface as compared to sham mice (Figure 1E and 1F, Supplemental Figure 2). For BDL treated animals, spironolactone normalized the apical intensity for each protein, decreasing it for the ENaC *γ* subunit and increasing it for AQP2. In contrast, spironolactone had no effect on the localization of either protein in sham mice. In contrast to these effects, spironolactone did not attenuate the effect of BDL on blood K^+^, suggesting this difference is independent of MR signaling (Figure 3, Supplemental Table 2).

**Figure 3.**
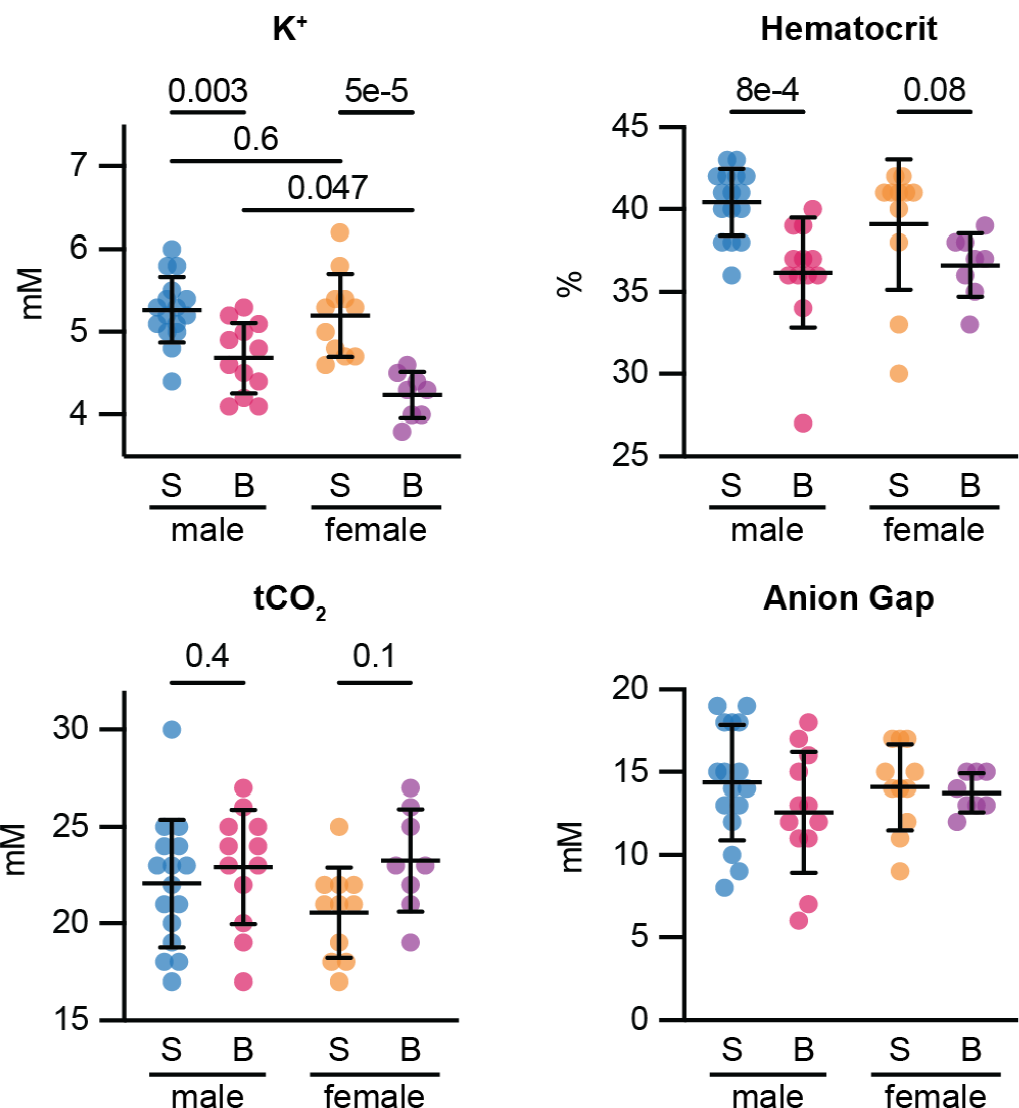
Effects of BDL with spironolactone treatment on blood electrolytes. Sham (S) and BDL (B) mice had chemistries from whole blood analyzed via an iSTAT analyzer while animals were anesthetized. Individual values with mean ± SD are shown for parameters that were affected by BDL (Table 1). Data were analyzed by two-way ANOVA with comparisons from post-hoc analysis shown. Summary statistics and two-way ANOVA results for all parameters are shown in Supplementary Table 2.

### Spironolactone reduces BDL-associated kidney injury

BDL alone induced apparent kidney injury within 11 days, however spironolactone treatment attenuated this damage. While spironolactone did not reduce histological evidence of injury (Figure 2A, B), it did reduce KIM-1 transcript levels in BDL mice by half, and NGAL transcript levels in BDL mice by 80% (Figure 2C). Spironolactone had no effect on KIM-1 or NGAL transcript levels in sham mice. In spironolactone treated female mice, BDL increased NGAL transcript levels over sham to a similar extent as spironolactone treated males (Supplemental Figure 1). BDL did not induce albuminuria, although spironolactone treatment had a modest general diminishing effect (Figure 2D). As spironolactone prevented the secondary effects of increased aldosterone after BDL, we included spironolactone in the remainder of our experiments to control for MR activation.

### BDL increases renal ENaC transport independent of MR

To measure the influence of BDL on ENaC-dependent renal transport, we determined how urine composition changed after administering benzamil, a diuretic that specifically blocks ENaC. Once recovered from surgery, all mice were given spironolactone and then housed in metabolic cages to facilitate 24-hour urine collections (Figure 4A). After collecting urine for a 24-hour basal period, we immediately administered benzamil (1.4 mg·kg^-1^·day^-1^ by intraperitoneal injection and simultaneously 1.4 mg·kg^-1^·day^-1^ by dietary supplement) and collected urine 24-hours later. Urine collected during both periods were analyzed and compared.

**Figure 4.**
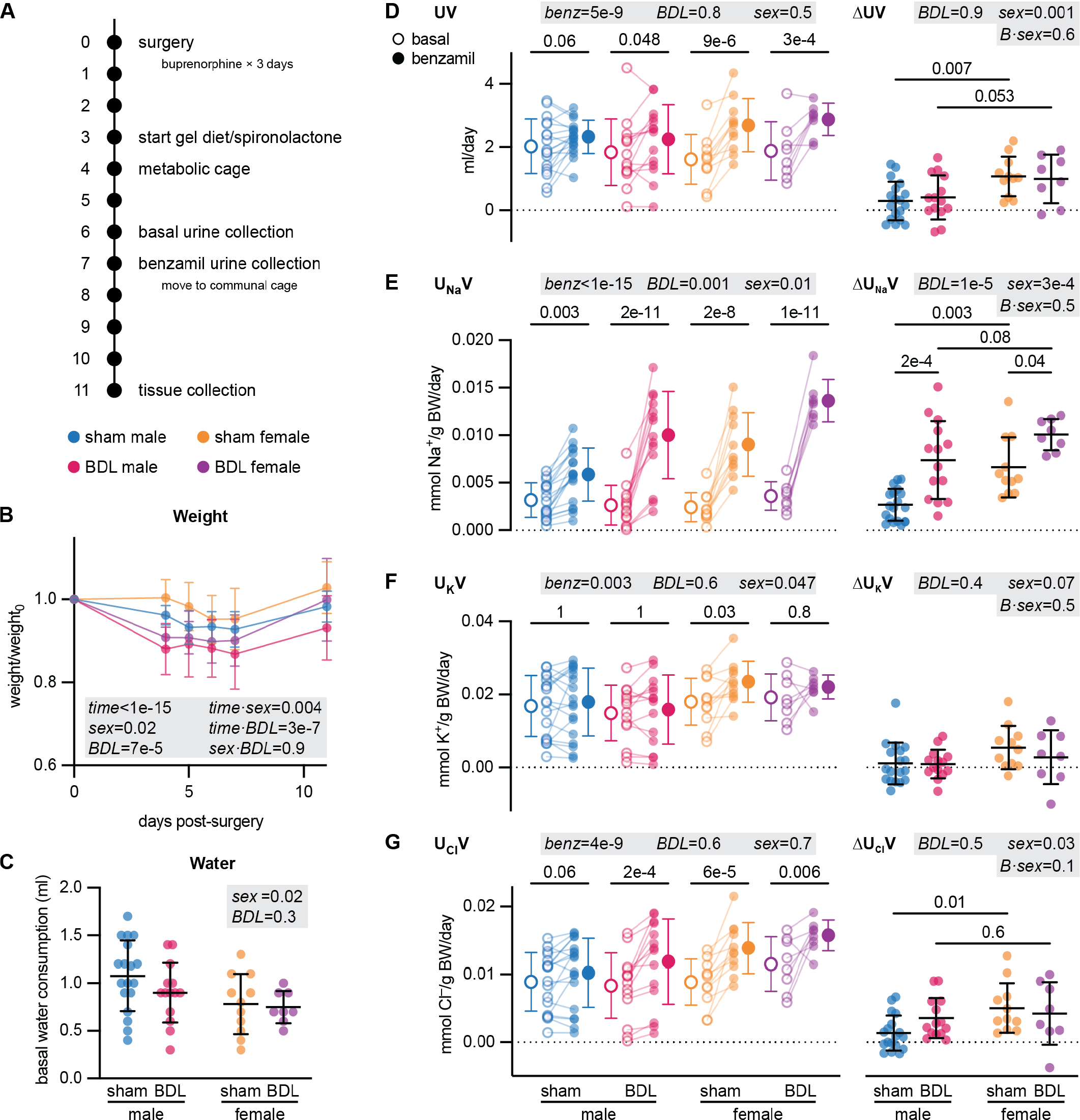
BDL with spironolactone treatment increased benzamil-sensitive natriuresis. *A*, Experimental timeline. *B*, Normalized animal weights (mean ± SD), analyzed by three-way ANOVA. *C*, Water consumption during the 24-hour basal urine collection period. *D-G*, Effect of benzamil on urinary volume (UV), Na^+^ (U_Na_V), K^+^ (U_K_V), and Cl^−^ (U_Cl_V). *Left*, 24-hour values before and after benzamil for individual animals, analyzed by repeated measures three-way ANOVA (shaded gray). All pairwise comparisons for U_Na_V are shown in Supplemental Table 3. *Right*, benzamil sensitive differences for each parameter (ΔU_X_V) were analyzed by two-way ANOVA (shaded gray). Individual values are shown for *C–G*.

Benzamil increased urinary Na^+^ excretion (U_Na_V) in each group (Figure 4E). Both BDL and sex were significant factors affecting benzamil-sensitive natriuresis (ΔU_Na_V). BDL increased ΔU_Na_V in males and females by 2.7- and 1.5-fold, respectively. These increases were similar to the ∼2-fold increases in ENaC open probability in split open mouse collecting ducts induced by 1 mM t-CA.^19^ In sham animals, females had 2.5-fold greater ΔU_Na_V than males, consistent with greater expression of active ENaC in females.^35^ For BDL mice, we did not detect differences in ΔU_Na_V between males and females.

Benzamil also increased urinary volumes (UV) and chloride (U_Cl_V) in most groups, although any changes were below the threshold of detection in sham males (Figure 4D, G). Sex was a significant factor for both benzamil-sensitive diuresis (ΔUV) and chloruresis (ΔU_Cl_V), and we detected differences between sham males and sham females for both parameters. Benzamil had little effect on urinary K^+^ secretion (U_K_V) in most groups in our experiments (Figure 4F), however, shorter collection time periods may be required to avoid compensation and detect differences instead of the 24-hour period we examined.^36^

During the experiment, mice transiently lost on average 5-13% of their initial body weight after surgery in each group, with weight recovery beginning after returning to communal cages (Figure 4A, B). Weight loss was greater for males compared to females, and for BDL mice compared to sham mice. During the basal 24-hour period, water consumption was greater for the larger males, but was not affected by BDL (Figure 4C).

### Bile acids regulate Na^+^ absorption in perfused isolated rabbit collecting ducts

To directly examine transport in renal tubules, we measured the effect of t-CA on ion fluxes in microperfused cortical collecting ducts dissected from rabbits. Like other conjugated primary bile acids, t-CA levels are greatly elevated after BDL in mice and in advanced liver diseases in humans.^16,17^ Tubules were sequentially perfused at 1 nL·min^-1^·mm^-1^ with control and 1 mM t-CA solutions, and the resulting ion fluxes were measured. Consistent with the ∼2-fold increase in ENaC currents observed in other systems,^18,19^ 1 mM t-CA increased net Na^+^ absorption by 2.6 ± 1.1-fold (Figure 5A). As collecting duct K^+^ secretion through apical K^+^ channels depends in large part on electrogenic ENaC-mediated Na^+^ reabsorption,^23^ we also examined effects on K^+^ secretion. In cortical collecting ducts perfused at slow flow rates, K^+^ secretion is mediated by the ROMK channel, whereas high flow rates trigger BK channel-mediated K^+^ secretion. t-CA had no effect on net K^+^ flux at low flow rates in our experiments. We then tested the effect of increasing the flow rate, which stimulates both Na^+^ reabsorption and K^+^ secretion in the absence of bile acids.^37,38^ In the presence of 1 mM t-CA, an increase in luminal flow further increased Na^+^ reabsorption by 2.2 ± 0.2-fold and K^+^ secretion by 3.5 ± 1.0-fold (Figure 5B). These data show that t-CA activates ENaC-mediated Na^+^ reabsorption but does not independently affect K^+^ secretion. They also show that flow-mediated effects on transport are preserved in the presence of t-CA.

**Figure 5.**
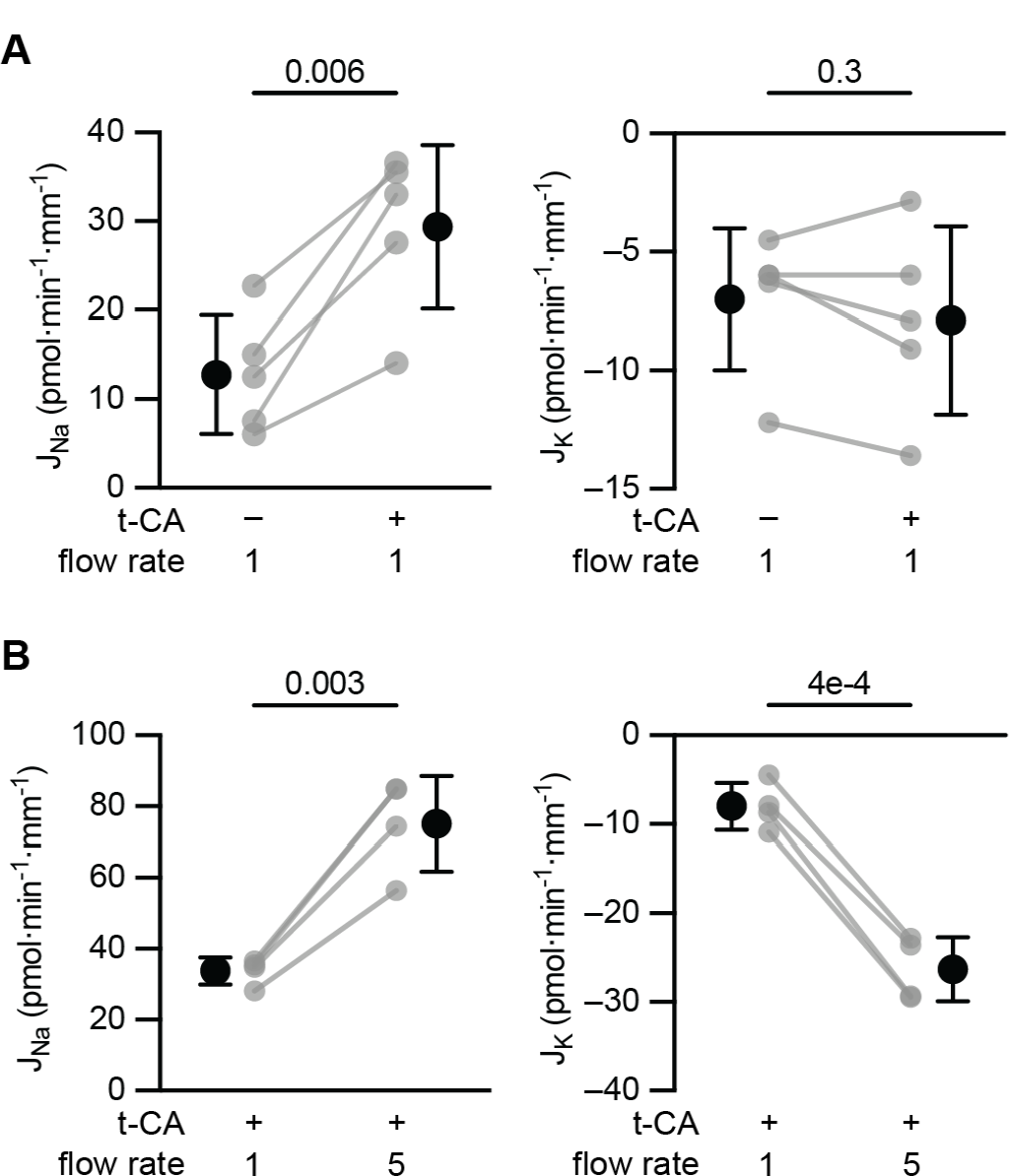
t-CA regulated Na^+^ flux in microperfused rabbit cortical collecting ducts isolated from rabbits. *A*, With a constant flow rate of 1 nL·min^-1^·mm^-1^, Na^+^ and K^+^ fluxes (*J*_Na_ and *J*_K_) were measured in the absence and then the presence of 1 mM t-CA. *B*, In the presence of 1 mM t-CA, *J*_Na_ and *J*_K_ was measured at 1 nL·min^-1^·mm^-1^, and then 5 nL·min^-1^·mm^-1^. Individual experiments (gray), mean and SD are shown for each experiment. Values were compared by paired Student’s *t* test.

### Na^+^ transporter abundance does not account for BDL effects on natriuresis

Since BDL could affect expression of transport proteins that influence urine composition, we examined their abundance in whole kidney lysates from our metabolic cage experimental cohort (Supplemental Figure 3). The Na-H-antiporter (NHE3), Na-K-2Cl-cotransporter (NKCC2), and Na-Cl-cotransporter (NCC) are upstream of ENaC in the nephron and their activity can influence Na^+^ delivery to ENaC. BDL reduced NHE3 expression in females, but we did not detect a reduction in males. BDL had no effect on NKCC2. BDL increased expression of total NCC in females but not males, but had no effect on the phosphorylated, active form (pNCC) in either sex. As shown above, BDL had no effect on ENaC α subunit expression after spironolactone treatment (Figure 1C-D, Supplemental Figure 1). For the β subunit, BDL had no effect in males, but reduced expression in females (Supplemental Figure 3). For the *γ* subunit, BDL increased expression of the full-length form in males but not females. We did not detect an effect on expression of the *γ* subunit’s cleaved form, associated with activation, in either females or males. We also detected no effect of BDL on AQP2 expression. Taken together, effects on ENaC expression were inconsistent between sexes and are unlikely to account for effects on urine composition.

### BDL drives fluid retention independent of MR

Because our animal model had increased benzamil-sensitive natriuresis, we hypothesized that these animals were experiencing fluid retention. We therefore measured body composition over a month using an EchoMRI^®^ machine. BDL mice transiently lost more weight after surgery but recovered to levels similar to sham mice (Figure 6A). This recovery in weight was largely due to an increase in total water, as this change composed a much larger proportion of the weight gain for BDL mice than for sham mice. We observed little increase in total water mass for sham mice over the month, whereas BDL mice gained nearly 2 g in total body water. Linear regression shows that BDL mice gained water at 9 times the rate of sham mice, so that total water accounted for 14 ± 18% of the weight gain for sham mice, but 70 ± 22% of the weight gain for BDL mice. As body water is a major component of measured lean mass, lean mass trends were similar as those seen for water, but any differences were below the threshold of detection. In contrast, fat mass does not contribute to total water mass and changed at similar rates for both sham and BDL mice.

**Figure 6.**
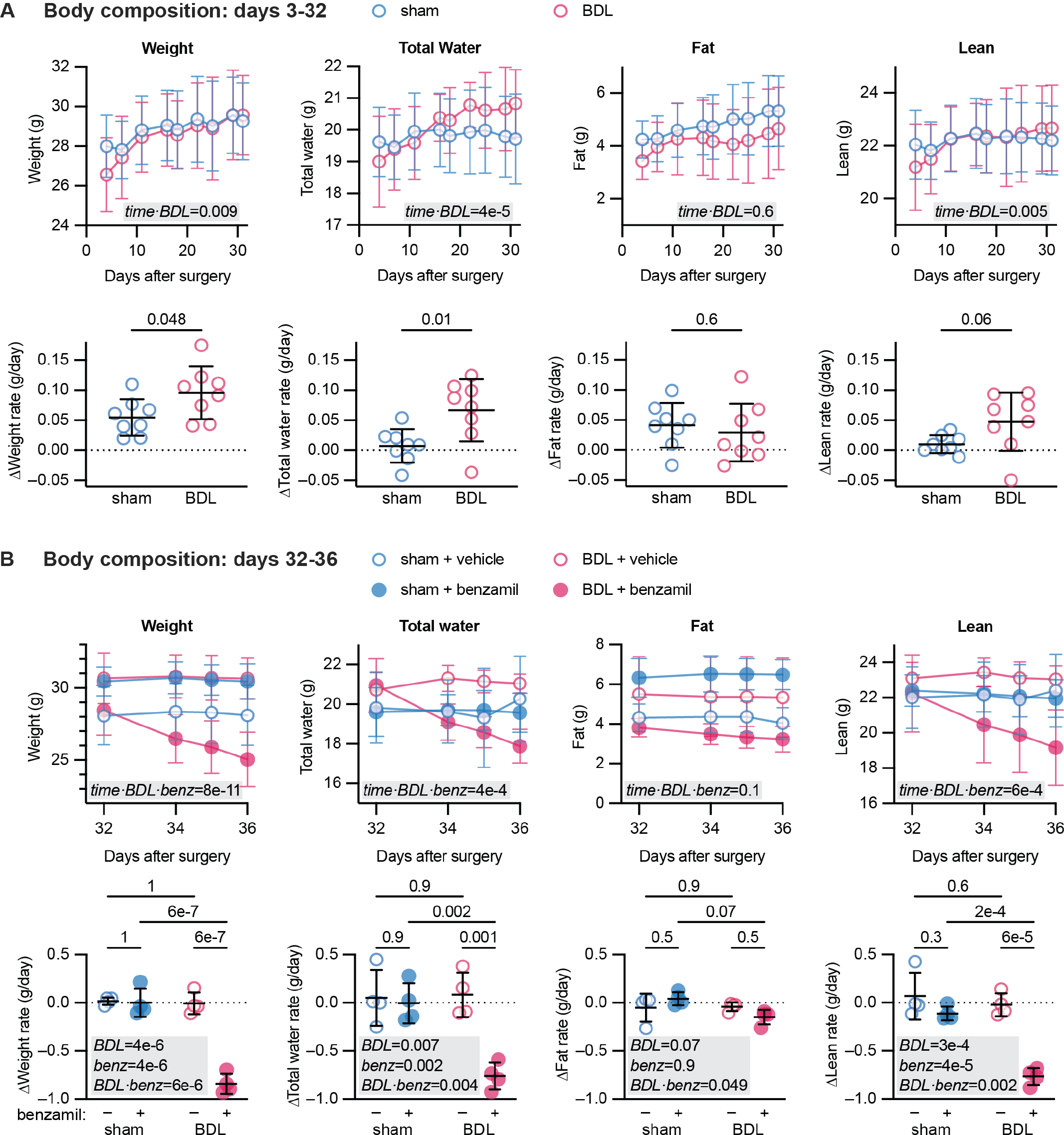
BDL promoted fluid retention and was reversed by benzamil. *A*, Three days after surgery, sham (*n*= 8) and BDL (*n* = 8) mice were started on spironolactone and body composition was measured over a month. Data were analyzed by repeated measures two-way ANOVA (interaction term shown, shaded gray) and linear regression to determine average rates of change. Rates of parameter change for each animal are plotted and were compared by Student’s *t* test. *B*, On days 32-36, half of each group (*n* = 4) was given benzamil or vehicle in the drinking water, and body composition measurements continued. Data were analyzed by repeated measures three-way ANOVA (full interaction term shown) and linear regression to determine average rates of change. Rates of parameter change for each animal are plotted and were compared by two-way ANOVA (shaded gray). Individual values are shown for fitted rates; mean and SD are shown for body composition measurements and rates of change.

To determine the contribution of ENaC-dependent transport to the fluid retention we observed, we then treated half of each group with benzamil while continuing spironolactone treatment for all groups (Figure 6B). Benzamil reduced weights in BDL mice (−0.8 ± 0.1 g/day) but not sham mice (0.0 ± 0.1 g/day). Strikingly, decreased total water mass in BDL mice (−0.8 ± 0.1 g/day) accounted for the entire weight reduction. Accordingly, we observed a similar decrease for lean mass but not fat mass. Collectively, these data show that BDL drove fluid retention and greatly enhanced benzamil sensitivity compared to sham in a model where the secondary effects of aldosterone activation have been eliminated.

## Discussion

Our results provide evidence for ENaC activation in liver disease independent of MR signaling. Our initial experiments demonstrate the effects of BDL on renal function through its ability to increase circulating bile acids and aldosterone (Figure 1). BDL increased expression of MR-regulated targets in the kidney (Figure 1) and increased renal damage (Figure 2). Using the MR antagonist spironolactone, we isolated the effects of BDL that were independent of secondary hyperaldosteronism. BDL increased benzamil-sensitive natriuresis independent of MR activation, suggesting higher ENaC activity after BDL (Figure 4E). BDL promoted fluid retention compared to sham control animals (Figure 6A), and blocking ENaC resulted in the loss of this excess water, again demonstrating ENaC’s involvement. Conversely, benzamil treatment did not impact total body water in sham animals, indicating minimal residual ENaC activity in sham mice when MR was inhibited (Figure 6B). Together, these data demonstrate increased ENaC activity independent of aldosterone signaling in a liver disease model with elevated plasma bile acids. The bile acid t-CA also increased net Na^+^ absorption by 2.6-fold in isolated microperfused cortical collecting ducts, ASDN segments that express apical ENaC. Notably, this technique enables transport to be examined in the absence of potentially confounding variables, such as paracrine factors and signaling. Several observations support the notion that higher levels of bile acids play a significant role in activating ENaC. When enteric delivery of bile acids is disrupted, bile acids are filtered and eliminated by the kidney where they come into contact with ENaC.^13,24^ Our isolated tubule experiments demonstrate that bile acids have direct effects on Na^+^ reabsorption (Figure 5). Additionally, even when we blocked MR signaling, BDL still increased ENaC activity, as measured by benzamil sensitivity. Except for expression of uncleaved ENaC γ in males, we observed no differences in ENaC expression, localization, or subunit cleavage that would account for increased ENaC activity in both males and females. This does not fit the classic paradigm of aldosterone regulation of ENaC, as aldosterone promotes increases in α subunit expression, channel localization at the apical surface, and cleavage of both the α and γ subunits. However, we and others have shown that bile acids activate membrane-resident channels instead of acting to increase the number of channels.^18-21^ These observations fit our current model more closely, as no increases in channel expression or localization are needed to account for the higher activity.

The etiology of fluid retention in liver disease is complex. It is widely thought that vasodilation and/or fluid sequestration reduces effective circulating volume to activate RAAS and renal Na^+^ retention. However, hemodynamic changes (low blood pressure) can either precede or follow the development of ascites,^39,40 41^ and fluid retention can occur without aldosterone elevation, suggesting alternative mechanisms.^2-11^ Indeed, renal Na^+^ retention occurs in cirrhotic animal models without increased aldosterone^42^ and precedes ascites formation.^41-44^ Even in hyperaldosteronism, aldosterone increases precede ascites formation, suggesting renal Na^+^ retention to be the cause of ascites, not the effect.^45^ We propose that bile acid activation of ENaC could be an independent mechanism for inappropriate Na^+^ avidity in liver disease.

Why do two separate mechanisms (aldosterone/MR and bile acids) ultimately affect the same channel in liver disease? For the former, liver dysfunction may slow aldosterone turnover and/or decrease sensed volume, leading to increased aldosterone release. Cirrhotic patients clear plasma aldosterone at half the rate of heathy subjects.^46^ The second mechanism of bile acids regulating ENaC may result from a physiologically advantageous interaction occurring in the wrong place. *In vitro*, bile acids directly bind ENaC and regulate its function.^19^ Binding occurs to at least two sites that are accessible from the luminal side with affinities of 50–250 μM. As ENaC is also expressed on the apical surface of cholangiocytes where luminal bile acids are in a similar range,^47,48^ we speculate that ENaC regulation by bile acids evolved as a mechanism to regulate the volume and composition of bile. Such a role has a been proposed for an ENaC paralog,^49^ and ENaC itself was proposed to regulate bile volume and composition through its response to flow.^47^ As disease shifts bile acid excretion to the kidney, we propose that this interaction becomes relevant in the nephron and promotes renal Na^+^ retention through ENaC.

Our study has important limitations. While we inhibited MR using spironolactone, aldosterone was still present at elevated levels and can stimulate ENaC apical expression.^50^ However, we did not observe increased ENaC γ subunit apical localization with BDL after spironolactone treatment. BDL will also increase renal tubular concentrations of other compounds, e.g., conjugated bilirubin, as tubular and urinary bilirubin precipitates suggest tubular concentrations can be high. We previously showed that bilirubin conjugates activate ENaC expressed in *Xenopus* oocytes,^18^ however, activation was modest compared to conjugated primary bile acids and required concentrations in excess of 1 mM. Decreased NHE3 expression in females after BDL may increase distal Na^+^ delivery and may account for some ENaC activation. However, the fact that BDL increased ENaC-mediated Na^+^ reabsorption in both sexes support additional mechanisms. BDL also increased expression of uncleaved ENaC *γ* in males, although it had no effect on expression of the cleaved form or apical localization. We cannot rule out contributions to our results from these effects.

Our findings suggest alternative strategies for managing fluid retention in liver patients. Initial treatment typically involves spironolactone.^1^ MR is constitutively activated by glucocorticoids in the late distal convoluted tubule and early connecting tubule, but is selectively activated by aldosterone in the late connecting tubule and collecting duct where it is protected from glucocorticoids by 11β-hydroxysteroid dehydrogenase type 2.^51^ As glucocorticoids are ∼1,000-fold more abundant than aldosterone, MR antagonists like spironolactone may poorly compete for MR in the early segments.^52^ In cases where plasma bile acids are elevated, a more effective approach might involve blocking ENaC using amiloride or triamterene. ENaC blockade could also complement the use of MR antagonists.^53^ In our experiments, ENaC blockade had a strong natriuretic and diuretic effect when administered, even with spironolactone present. Another potential approach is to target the reduction of circulating bile acids. Currently available bile acid sequestrants (e.g., cholestyramine) depend on bile acid delivery to the gut, which may be impaired in liver disease. Therefore, compounds that specifically target metabolic or transport processes to reduce plasma bile acids may also be a promising approach.

In summary, our work provides evidence supporting a novel mechanism for ENaC activation in a cholestatic liver disease model: direct activation by luminal bile acids. This mechanism does not depend on MR signaling. Activation increased renal Na^+^ retention through ENaC and decreased blood K^+^ levels. It also enhanced fluid retention, which was highly responsive to ENaC blockers. In isolated tubules, bile acids enhanced Na^+^ absorption, but did not affect K^+^ secretion or flow-dependent K^+^ secretion. These data suggest that any bile acid effect on K^+^ secretion is secondary to effects on ENaC transport. We conclude that this mechanism likely contributes to fluid retention in conditions where plasma bile acid levels are elevated, and that it should be considered in the management of patients.

## Abbreviations

ENaC: epithelial Na^+^ channel
RAAS: renin-angiotensin-aldosterone system
MR: mineralocorticoid receptor
BDL: bile duct ligation
t-CA: taurocholic acid
AQP2: aquaporin 2
KIM-1: Kidney Injury Molecule-1
NGAL: neutrophil gelatinase-associated lipocalin
ATP6V1B1: B1 subunit of the vacuolar-ATPase proton pump

## Data Availability

All data are available in the manuscript and supplementary materials.

## Authors’ Contributions

Conceptualization: XW, SMM, OBK

Methodology: XW, SMM, RC, CJB, MJJ, LMS, OBK

Investigation: all

Writing – Original Draft: OBK

Writing – Review & Editing: XW, SMM, RC, CJB, RJT, MJJ, LMS, OBK

Funding acquisition: RJT, MJJ, OBK

## Acknowledgments

We thank Ramon Bataller, Carlos Fernández Carrillo, Andrew Nickerson, Evan Ray, Ora Weisz, and Thomas Kleyman for helpful discussions.

## Notes

Conflict of Interest: Authors declare no conflict of interest.

Financial Support: This work was supported by National Institute of Health Grants R01 DK125439, S10 OD021627, P30 DK120531, P30 DK079307, R03 DK131093, I01 BX005680, R01 DK131991, T32 DK061296, T32 DK007052 and the Pittsburgh Foundation (MR2020 109502).

### Competing Interest Statement

The authors have declared no competing interest.

