## Supplemental Materials for "Mineralocorticoid receptor-independent activation of ENaC in bile duct ligated mice"

**Supplementary Methods**

*Materials*

Animal protocols conformed with the National Institutes of Health Guide for the Care and Use of Laboratory Animals and were approved by the Institutional Animal Care and Use Committees of the University of Pittsburgh and the Icahn School of Medicine at Mount Sinai. Eight- to twelve-week-old 129-Elite mice were purchased from Charles River Laboratories (Wilmington, MA) and housed in a temperature-controlled facility at the University of Pittsburgh on a standard 12-hour light/dark cycle. Adult (>6 wk) female New Zealand White rabbits, purchased from Charles River, were housed in the Center for Comparative Medicine at the Icahn School of Medicine at Mount Sinai. All animals were given free access to water and chow. Benzamil was a gift from Dr. Thomas Kleyman. Spironolactone and t-CA were purchased from Thermo Fisher Scientific and Cayman chemical, respectively.

*Animal model: BDL*

BDL was used to increase plasma bile acids as previously shown.[1] Male and female 129-Elite mice approximately 8-12 weeks of age were subjected to sham or common BDL operation, as previously described.[2] Briefly, animals were anesthetized with isoflurane and shaved of fur on the abdominal skin. A midline laparotomy of 1-2 cm was made to open the top of the peritoneal cavity. The liver was shifted away from the midline with a cotton swab soaked in saline, and the gut was moved down to expose the bile duct. The bile duct was carefully separated from the portal vein and hepatic artery using forceps and 5-0 suture was used to completely ligate the duct in two separate locations, however, the duct was not cut between these sutures. Both the muscle and skin were sutured closed, buprenorphine was administered to manage pain, and the mice were monitored until fully recovered.

*Metabolic cages experiments*

Spironolactone treatment (20 $mg \cdot kg^{-1} \cdot day^{-1}$ via drinking water) was initiated 3 days after surgery. Mice were then transferred to metabolic cages (Tecniplast) to facilitate 24-hour urine collections. They were acclimated to gel diet composed of regular mouse chow, ground to a fine powder, mixed with 1-2% agar and water. On day 6, 24-hour baseline urine samples were collected, followed by benzamil administration (1.4 $mg \cdot kg^{-1} \cdot day^{-1}$ intraperitoneal injection and 1.4 $mg \cdot kg^{-1} \cdot day^{-1}$ added to the gel diet). On day 7, the resulting 24-hour urine was collected. Urinary $Na^+$, $K^+$ and $Cl^-$ concentrations were measured using an EasyLyte Plus ion-specific electrode system (Medica). Benzamil-sensitive changes were assessed as the difference between day 6 and day 7 urinary values. On Day 11, mice were anesthetized with isoflurane and blood was collected with a heparinized syringe via cardiac puncture and immediately analyzed using an iSTAT (Abbot). Kidneys were harvested and flash frozen for immunoblot analysis or fixed in either 10% formalin for IHC or 4% paraformaldehyde for immunofluorescence.

*Body Composition*

Spironolactone treatment (20 $mg \cdot kg^{-1} \cdot day^{-1}$ via drinking water) was started 3 days after BDL surgery, and body composition was measured using an EchoMRI™-100H (EchoMRI LLC). The EchoMRI™ measures body composition using NMR spectroscopy and the resonant frequency of hydrogen and the differences in relaxation times of hydrogen proton spins in different physiological environments, allowing quantification of total lean, fat and body water mass.[3] Benzamil (1.4 $mg \cdot kg^{-1} \cdot day^{-1}$) or vehicle (dimethylsulfoxide) was added to drinking water, as indicated.

*Measurement of plasma and urinary compounds*

Plasma aldosterone and total bile acid levels were measured using an aldosterone ELISA kit (Enzo Life Sciences, ADI-900-173) and bile acid assay kit (Sigma, MAK309), respectively, according to the manufacturer's protocols. Samples were diluted 18-fold in assay buffer or 5-fold in water prior to measure plasma aldosterone or total bile acids.

Urine albumin levels were measured using the ELISA Starter Accessory kit (Bethyl Laboratories, E101), according to the manufacturer's protocols. Briefly, plates were coated with goat anti–mouse albumin antibody (Bethyl Laboratories, E90-134A). Serially diluted mouse reference serum (ICL Lab, RS-90AL) and urine samples diluted 1000-fold in water were incubated in the plate for 1 h at room temperature, followed by washes. HRP-conjugated secondary antibody (Bethyl Laboratories, A90-134P, 1:80,000) was applied for 1 h at room temperature. Following washes, 3,3',5,5'-Tetramethylbenzidine was added and color development was terminated by addition of 0.2 M $H_2SO_4$. Absorbance was measured at 450 nm in a microplate reader. The standard curve was fitted using linear/sigmoidal fit in GraphPad Prism. Albuminuria is reported as albumin concentration normalized to 24-hour urinary volume.

*Quantitative Reverse Transcriptase PCR*

Kidney samples were homogenized with TRIzol Reagent (Invitrogen) and total RNA was isolated. cDNA was generated with the RevertAid Reverse Transcriptase Kit (ThermoFisher Scientific). Quantitative PCR was performed using iTAQ™ Universal SYBR Green Supermix (Bio-Rad) on the CFX Connect™ Real-Time System (Bio-Rad Laboratories). Each reaction contained one of the following primer pairs: mouse *β-Actin* (Forward: 5′- CGCAGCCACTGTCGAGTC-3′; Reverse: 5′-GTCATCCATGGCGAACTGGT-3′), mouse *Kim1* (Forward: 5′-GGAATCCCATCCCATACTCCT-3′; Reverse: 5′-AAGTATGTACCTGGTGATAGCCAC-3′), mouse *Ngal* (Forward: 5′-CCATCTATGAGCTACAAGAGAACAAT-3′; Reverse: 5′-TCTGATCCAGTAGCGACAGC-3′). Results were normalized to *β-Actin* and gene expression was determined with the $2^{-\Delta\Delta Ct}$ method.[4]

*Immunoblotting*

Kidneys were homogenized in 200 μL of ice-cold commercial RIPA buffer containing 50 mM Tris-Cl, 5 mM EDTA and 1% Triton X-100 (pH 7.5) supplemented with Cocktail protease inhibitor (Calbiochem) and PhosSTOP phosphatase inhibitor (Roche Diagnostics). Homogenates were centrifuged at 12,000 rpm for 10 min at 4°C and sediment was discarded. Protein concentration was determined using the BCA colorimetric assay according to the manufacturer's instructions (Pierce Rapid Gold BCA Protein Assay Kit, Thermo Fisher Scientific). Samples were prepared in 4x Laemmli buffer (Biorad) and were incubated at room temperate for 10min, 60 °C for 6 min, or 97 °C for 3 min, depending on the protein target. Samples (60 μg/lane) were separated on 4-15% or 8-16% Criterion TGX stain-free gels (Biorad) at 150 V for 90 min and transferred to a nitrocellulose membrane for 40 minutes at 400 mA. Nitrocellulose membranes were blocked with 5% non-fat milk in PBS and incubated with primary antibodies as indicated in Supplemental Table 1 overnight at 4 °C. After 3 washes with PBS, the membrane was incubated with HRP-conjugated goat anti-rabbit (1:10000; Jackson ImmunoResearch Laboratories) secondary antibodies as indicated for 1 hour at room temperature. Blots were developed using Clarity™ Western Blotting Substrate, followed by ClarityMax when needed (BioRad). Blots were imaged on a ChemiDoc™ Imaging System (BioRad). Blot quantification was performed using densitometry in Image lab (BioRad).

*Indirect immunofluorescence microscopy and quantification*

Formalin fixed kidneys were dehydrated and embedded in paraffin blocks, which were then sectioned at 3 μm onto charged slides. Slides were rehydrated in xylene followed by graded ethanol concentrations, and antigen retrieval was performed in citrate buffer. Kidney slices were immunolabeled with an antibody against ENaC-$\gamma$ raised in rabbit at 1:200 dilution in Dako low background diluent (Dako North America, Carpinteria, CA), followed by a secondary antibody raised in the donkey coupled to CY3 (1:800 dilution; Jackson ImmunoResearch Laboratories, Westgrove, PA). Slices were also colabeled with antibody raised in the goat against water channel aquaporin-2 (AQP2; Santa Cruz Biotechnology, Dallas, TX) at 1:400 dilution followed by secondary antibody raised in the donkey coupled to Alexa 488 (1:400 dilution; Jackson ImmunoResearch Laboratories, Westgrove, PA). Immunolabeled tissues were mounted in SlowFade Glass Antifade mounting medium (ThermoFisher, Grand Island, NY).

Slides were blinded before being imaged on an Leica SP8 inverted confocal microscope as follows: overviews (1X zoom) of kidney cortex and for quantitation, four images per cortical region (superficial cortex, midcortex and juxtamedullary) from each slide at 4X zoom using a 40x oil 1.3 N.A objective and step size of 0.35 micron. Zoomed images were selected to optimize number of tubules with open lumens for subsequent analysis. Conventional settings for Alexa Fluor 488 and Cy3 were used with laser lines at 488 and 552 nm, hybrid detectors and sequential scanning.

While blinded, maximum projection images were analyzed using LAS X 3.5.519976, version 15 (Leica). Background values for the lumen was estimated at 0.9% for ENaC and 0.5% for AQP2 and used for subsequent processing of all groups. ENaC and AQP2 positive tubules were identified, and line scans were drawn across individual cells starting from the lumen. ENaC and AQP2 signal profiles were then analyzed using Matlab R2023a (Mathworks, Inc.). For each pair of profiles, the apical edge was defined as the first value above the lumen background for either signal. Traces whose apical minima were above lumen background were discarded. The common basal edge was defined as the minimum value beyond the peak identified in the basal region (i.e., final third) of the line trace. Signal profiles were trimmed to the common defined peaks, and cumulative intensity for the apical 2 μm of each scan was calculated. Profiles and cumulative intensity were visualized using Prism 9 (GraphPad Software, LLC).

*Isolation, perfusion, and measurement of ion fluxes in rabbit cortical collecting ducts*

Kidneys were removed via a midline incision, and single tubules dissected freehand in cold (4 °C) $Na^+$-Ringer solution containing (in mM): 135 NaCl, 2.5 $K_2HPO_4$, 2.0 $CaCl_2$, 1.2 $MgSO_4$, 4.0 lactate, 6.0 L-alanine, 5.0 HEPES, and 5.5 D-glucose, pH 7.4, 290 ± 2 mOsm/kg, as previously described.[5] A single tubule was isolated from each animal and immediately transferred to a temperature- and $O_2$/$CO_2$-controlled specimen chamber set on the stage of a Nikon inverted epifluorescence microscope (Eclipse Ti), mounted on concentric glass pipettes, and microperfused and bathed at 37 °C with Burg's solution containing (in mM): 120 NaCl, 25 $NaHCO_3$, 2.5 $K_2HPO_4$, 2.0 $CaCl_2$, 1.2 $MgSO_4$, 4.0 $Na^+$ acetate, 1.0 $Na_3$ citrate, 6.0 L-alanine, and 5.5 D-glucose, pH 7.4, 290±2 mOsm/kg.[5] During the 45 min equilibration period and thereafter, the perfusion chamber was continuously suffused with a gas mixture of 95% $O_2$-5% $CO_2$ to maintain pH at 7.4 and 37 °C. The bathing solution was continuously exchanged at a rate of 10 ml/hr using a syringe pump (Razel, Stamford, CT).

Transport measurements were performed in the absence of transepithelial osmotic gradients and thus water transport was assumed to be zero. Three to four samples of tubular fluid were collected under water-saturated light mineral oil by timed filling of a calibrated ~7 nl volumetric constriction pipette at slow (~1) and fast (~5 $nl \cdot min^{-1} \cdot mm^{-1}$) flow rates, in the absence or presence of luminal t-CA (1 mM), as indicated. To determine the concentrations of $K^+$ and $Na^+$ delivered to the tubular lumen, ouabain (200 μM) was added to the bath at the conclusion of each experiment to inhibit all active transport, and an additional three to four samples of tubular fluid were obtained for analysis.[6,7]

The cation concentrations of perfusate and collected tubular fluid samples were determined by helium glow photometry and the rates of net transport ($J_x$, in $pmol \cdot min^{-1} \cdot mm^{-1}$ tubular length) were calculated using standard flux equations, as previously described.[8] The calculated ion fluxes were averaged to obtain a single mean rate of ion transport for the CCD at each flow rate under each condition. The flow rate was varied by adjusting the height of the perfusate reservoir.

*Statistical analysis*

The results of all individual experiments are presented in the figures, where practicable. All summary statistics are mean ± SD, except where noted. Statistical tests used are indicated and were performed using Prism 9.5.0. For pair comparisons, normality was first tested using the Shapiro-Wilk test. Normally distributed data were then analyzed using Student's *t* test and non-normally distributed data were analyzed using the Mann-Whitney U test. Data with two or three factors were analyzed using two-way ANOVA or three-way ANOVA tests, respectively. Repeated measures versions of each test were used as indicated. Post-hoc pairwise comparisons were performed using Holm-Šidák's multiple comparison test. Where possible, multiple comparisons were limited to factors that the overall test indicated were sources of variation. P-values below 0.05 were considered significant. All p-values are shown with 1 significant figure, except p-values that rounded to 0.05, where 2 significant figures are shown.

| Antibody | Source | Catalog number | Dilution for blot |
|---|---|---|---|
| ENaC $\alpha$ subunit | Gift from Johanne Loffing | | 1:2000 |
| ENaC $\beta$ subunit | StressMarq | SPC-404 | 1:1000 |
| ENaC $\gamma$ subunit | StressMarq | SPC-405 | 1:1500 |
| NCC | Millipore-Sigma | AB3553 | 1:1000 |
| pNCC | Phosphosolutions | P1311-53 | 1:1000 |
| NKCC2 | StressMarq | SPC-401D | 1:1000 |
| NHE3 | StressMarq | SPC-400D | 1:1000 |
| SGK1 | Cell signaling technology | 12103 | 1:1000 |
| AQP2 | Santa Cruz | SC-515770 | 1:1000 |
| ATP6V1B1 | BiCell Scientific | 20901 | 1:1000 |

**Supplemental Table 1. Antibodies used**

| | **sham ♂ (16)** | **BDL ♂ (12)** | **sham ♀ (11)** | **BDL ♀ (8)** | $\boldsymbol{P}_{\text{BDL}}$ | $\boldsymbol{P}_{\text{sex}}$ |
|---|---|---|---|---|---|---|
| $Na^+$ (mM) | 144 ± 3 | 144 ± 3 | 144 ± 1 | 145 ± 2 | 0.2 | 0.6 |
| * $K^+$ (mM) | 5.3 ± 0.4 | 4.7 ± 0.4 | 5.2 ± 0.5 | 4.2 ± 0.3 | **2e-7** | **0.046** |
| $Cl^-$ (mM) | 114 ± 3 | 115 ± 3 | 115 ± 4 | 113 ± 3 | 0.7 | 0.8 |
| $iCa^{2+}$ (mM) | 1.22 ± 0.07 | 1.18 ± 0.13 | 1.20 ± 0.18 | 1.22 ± 0.05 | 0.8 | 0.5 |
| * $tCO_2$ (mM) | 22 ± 3 | 23 ± 3 | 21 ± 2 | 23 ± 3 | **0.049** | 0.5 |
| * AnGap (mM) | 14 ± 3 | 13 ± 4 | 14 ± 3 | 14 ± 1 | 0.3 | 0.6 |
| BUN (mg/dL) | 27 ± 3 | 25 ± 5 | 24 ± 3 | 26 ± 3 | 0.9 | 0.5 |
| * Hct (%) | 40 ± 2 | 36 ± 3 | 39 ± 4 | 37 ± 2 | **4e-4** | 0.6 |

**Supplemental Table 2. Effects of BDL with spironolactone treatment on blood electrolytes.** Blood was collected 11 days after surgery. Groups were compared by two-way ANOVA, with p-values for each factor shown. * Data and multiple comparisons are detailed in Figure 3.

| | female BDL benzamil | female BDL basal | female sham benzamil | female sham basal | male BDL benzamil | male BDL basal | male sham benzamil |
|---|---|---|---|---|---|---|---|
| female BDL basal | **1e-11** | | | | | | |
| female sham benzamil | **0.006** | 7e-4 | | | | | |
| female sham basal | 2e-12 | **1** | **2e-8** | | | | |
| male BDL benzamil | **0.03** | 1e-5 | 1 | 1e-8 | | | |
| male BDL basal | 5e-13 | **1** | 2e-6 | 1 | **2e-11** | | |
| male sham benzamil | 3e-8 | 0.4 | **0.03** | 0.02 | **8e-4** | 0.02 | |
| male sham basal | 7e-13 | 1 | 4e-6 | **1** | 9e-9 | **1** | **0.003** |

**Supplemental Table 3**. Repeated measures three-way ANOVA with Holms-Šidák multiple comparisons for $U_{Na}V$ response to benzamil. Sex (p=0.01), BDL (p=0.001), and benzamil (p<1e-15) were all sources of variation. Benzamil interacted with sex (p=3e-4) and BDL (p=1e-5), but sex did not interact with BDL (p=0.4). Notes: all groups responded to benzamil. No differences between basal UNaV for any groups. BDL + benzamil was higher than BDL-basal for both males and females.

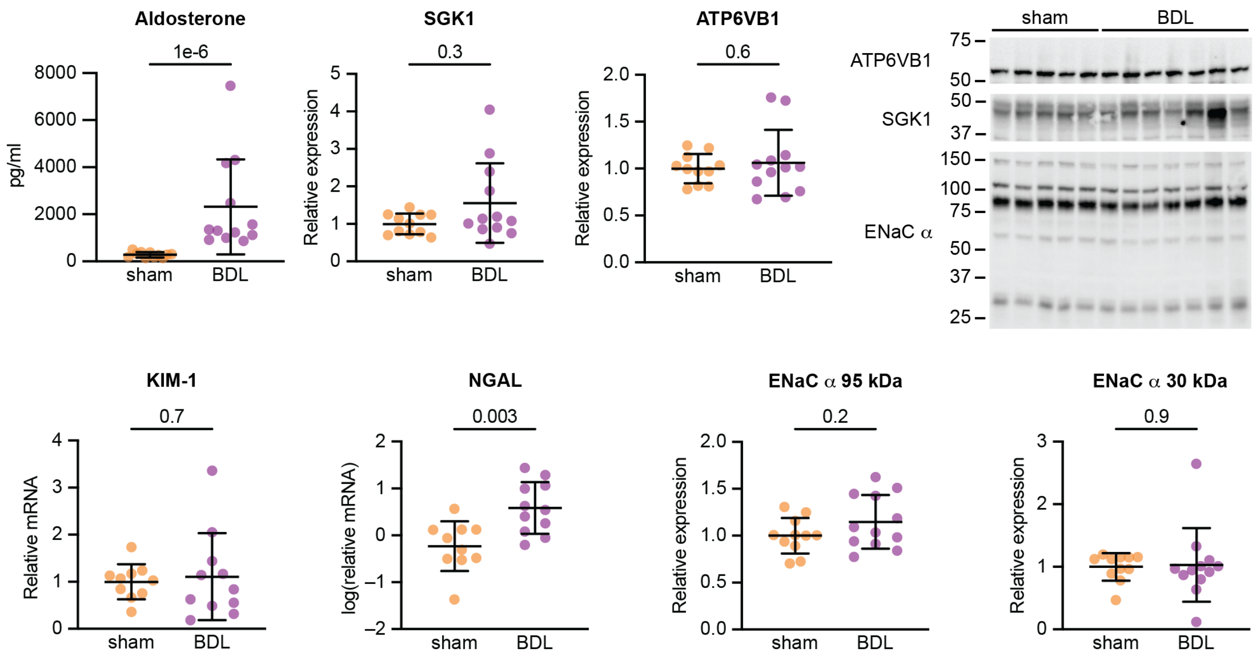


**Supplemental Figure 1. Effect of BDL with spironolactone in female mice.** Plasma aldosterone was compared by Mann-Whitney U test. Transcript levels of kidney injury markers NGAL and KIM-1 were measured by qPCR from kidney homogenates. SGK1, ATP6VB1, and ENaC α protein expression were evaluated by immunoblot from kidney homogenates. Values were normalized to sham, and individual points, mean, and SD are shown for each. Expression was compared by Student's *t* test.

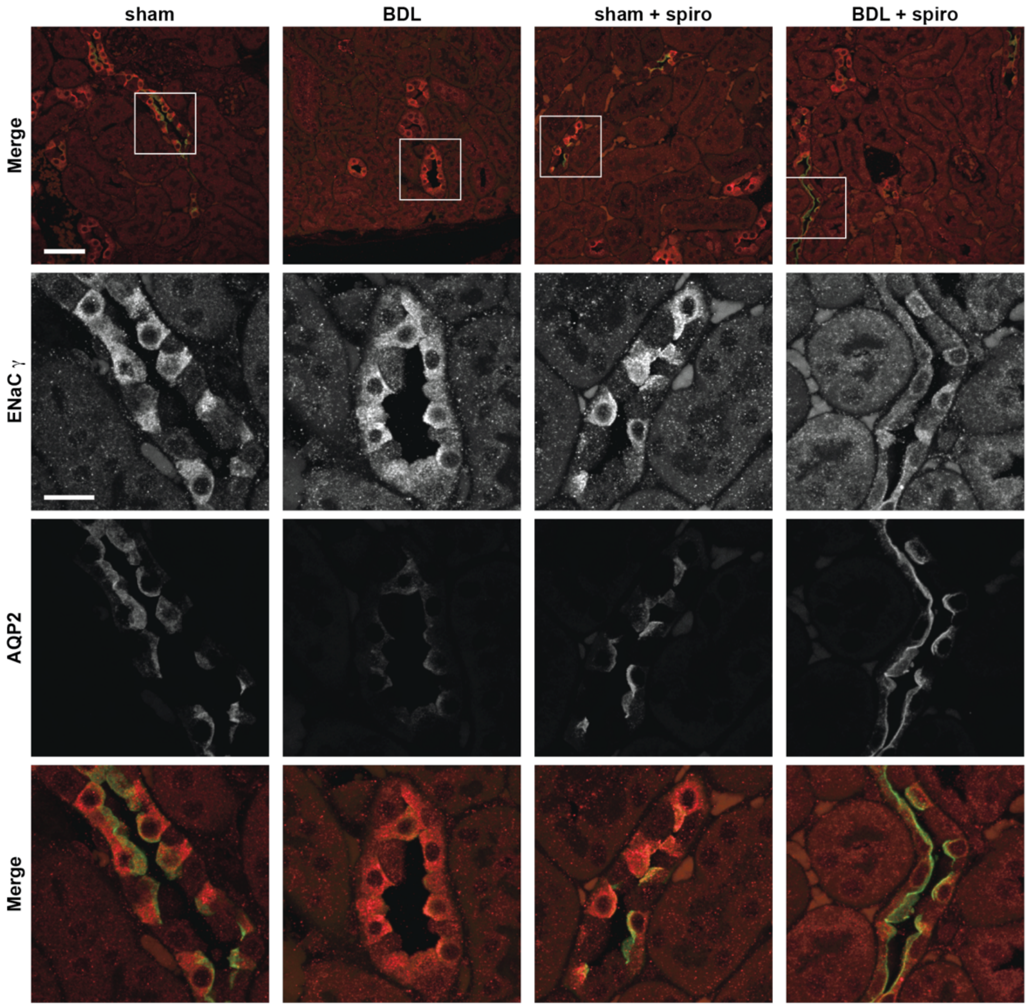


**Supplemental Figure 2. Spironolactone normalizes BDL effects on ENaC and AQP2 apical localization.** Representative maximum projection images of kidney cortex labeled with antibodies directed against the ENaC γ subunit (red) and AQP2 (green). Scale bar is 50 µm for the top row and 15 µm for the remainder. AQP2-positive tubules were selected for quantification.

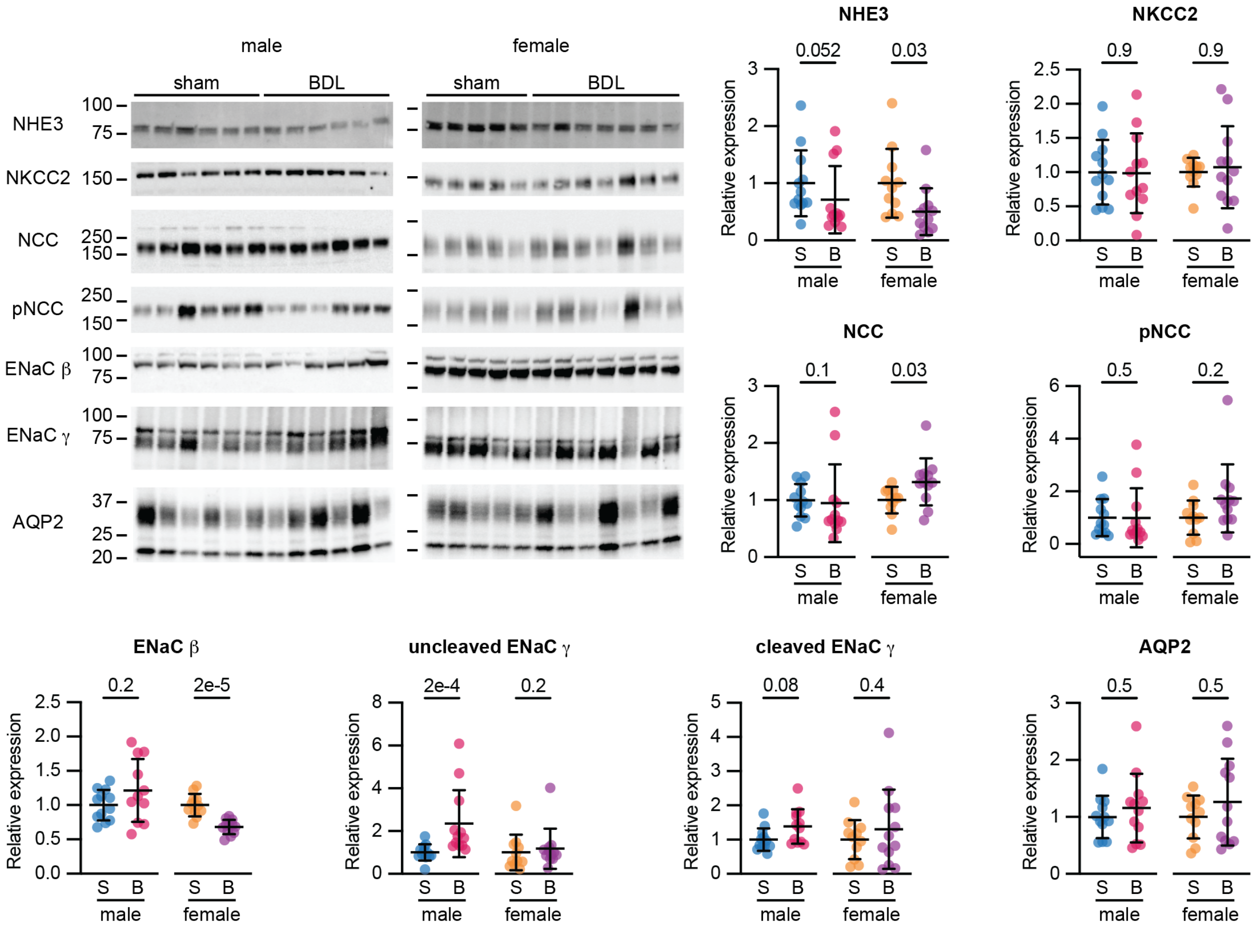


**Supplemental Figure 3. Effect of BDL on expression in the presence of spironolactone.**

Transporter expression in kidney homogenates was evaluated by immunoblot and normalized to sham. Individual points with mean and SD shown. Normally distributed data (ENaCs and AQP2) were compared by Student's *t* test. Other groups were compared Mann-Whitney U test.
